# Randomly incorporated genomic 6mA delays zygotic transcription initiation

**DOI:** 10.1101/2022.10.27.514014

**Authors:** Febrimarsa, Sebastian G Gornik, Sofia N Barreira, Miguel Salinas-Saavedra, Christine E Schnitzler, Andreas D Baxevanis, Uri Frank

## Abstract

N6-methyldeoxyadenosine (6mA) is a chemical alteration of DNA, observed across all realms of life. The functions of 6mA are well understood in bacteria but its roles in animal genomes have been controversial. We show that 6mA randomly accumulates in early embryos of the cnidarian *Hydractinia symbiolongicarpus*, with a peak at the 16-cell stage followed by clearance to background levels two cell cycles later, at the 64-cell stage – the embryonic stage at which zygotic genome activation occurs in this animal. Knocking down *Alkbh1*, a putative initiator of animal 6mA clearance, resulted in higher levels of 6mA at the 64-cell stage and a delay in the commencement of zygotic transcription. Our data are consistent with 6mA originating from recycled nucleotides of degraded m6A-marked maternal RNA post-fertilization. Therefore, while 6mA does not function as an epigenetic mark in *Hydractinia*, its random incorporation into the early embryonic genome inhibits transcription. Alkbh1 functions as a genomic 6mA ‘cleaner’, facilitating timely zygotic genome activation. Given the random nature of genomic 6mA accumulation and its ability to interfere with gene expression, defects in 6mA clearance may represent a hitherto unknown cause of various pathologies.

## INTRODUCTION

Methylation of adenine in DNA (6mA) and the functions it fulfils are well documented in bacteria (Geier and Modrich, 1979; Haagmans and van Der Woude, 2000; Lahue et al., 1987; Slater et al., 1995) and protists (Beh et al., 2019; Wang et al., 2019), but studies on this DNA modification in animals have revealed conflicting reports (Bochtler and Fernandes, 2020; Douvlataniotis et al., 2020; Kong et al., 2022). Low levels of 6mA were reported in the genomes of flies (He et al., 2019; Yao et al., 2018; Zhang et al., 2015), worms (Greer et al., 2015; O’Brown et al., 2019), fish (Liu et al., 2016; O’Brown *et al*., 2019), and mammalian cells (Koziol et al., 2016; Wu et al., 2016; Xiao et al., 2018; Xie et al., 2018), and were shown to correlate with transposon transcripts level in flies and mouse cells (Wu *et al*., 2016; Xie *et al*., 2018; Zhang *et al*., 2015). However, some of these studies were challenged by others, attributing their findings to antibody artifacts (Abakir et al., 2020; Douvlataniotis *et al*., 2020) or to bacterial contamination (Kong *et al*., 2022; O’Brown *et al*., 2019; Schiffers et al., 2017). To address this apparent discrepancy, we have studied 6mA during early embryogenesis of *Hydractinia symbiolongicarpus*, a member of the early-diverging phylum Cnidaria. As a sister group to Bilateria, cnidarians may provide new insights into the evolution of animal traits. We report a peak in the level of 6mA in 16-cell stage embryos. However, 6mA marks were randomly distributed in the genome, inconsistent with having an epigenetic function. We find that the clearance of 6mA before the 64-cell stage by the dioxygenase Alkbh1 is necessary for timely zygotic genome activation (ZGA). We propose that 6mA is passively and randomly accumulated in the genome due to the rapid degradation of m6A-marked maternal RNA, NTP-dNTP conversion by ribonucleotide reductase, and random integration into the early embryonic genome.

## RESULTS

### Dynamics and distribution of 6mA during embryogenesis

To quantitatively assess 6mA levels in *Hydractinia*, we extracted genomic DNA from adult specimens and from different embryonic stages. The samples were then enzymatically digested and purified. Synthetic oligonucleotides containing 6mA were similarly treated and used as external standards for ultra-high-performance liquid chromatography coupled with triple-quadrupole tandem mass spectrometry (UHPLC-QQQ) (Figure 1A). We found that the levels of 6mA were at background level in sperm (~0.015% 6mA/dA mol/mol) and slightly above background at the two-cell stage, but increased gradually to ~0.06% in 16-cell stage embryos. Levels of 6mA rapidly decreased to background level by the 64-cell stage and were maintained at this level to adulthood, being indistinguishable from the negative control (Figure 1B). We re-analyzed the level of 6mA/dA in 16- and 64-cell stage embryos by ultra-high performance liquid chromatography coupled with quadrupole ion trap tandem-mass spectrometry (UHPLC-QTRAP) using stable isotope-labeled [^3^D1]-6mA as an internal standard for sample enrichment and quantitation (Figure 1A). This method enabled us to detect 6mA/A levels of 0.01% as being distinct from the negative control (Figure 1C and S1A) and confirmed the 6mA data at the 16- and 64-cell stages obtained by HPLC-QQQ (Figure 1B-C). Hence, 6mA levels are dynamic in early embryos, being low in early embryos, high at the 16-cell stage, and low again at the 64-cell stage and later (Figure 1B-C).

**Figure 1.**
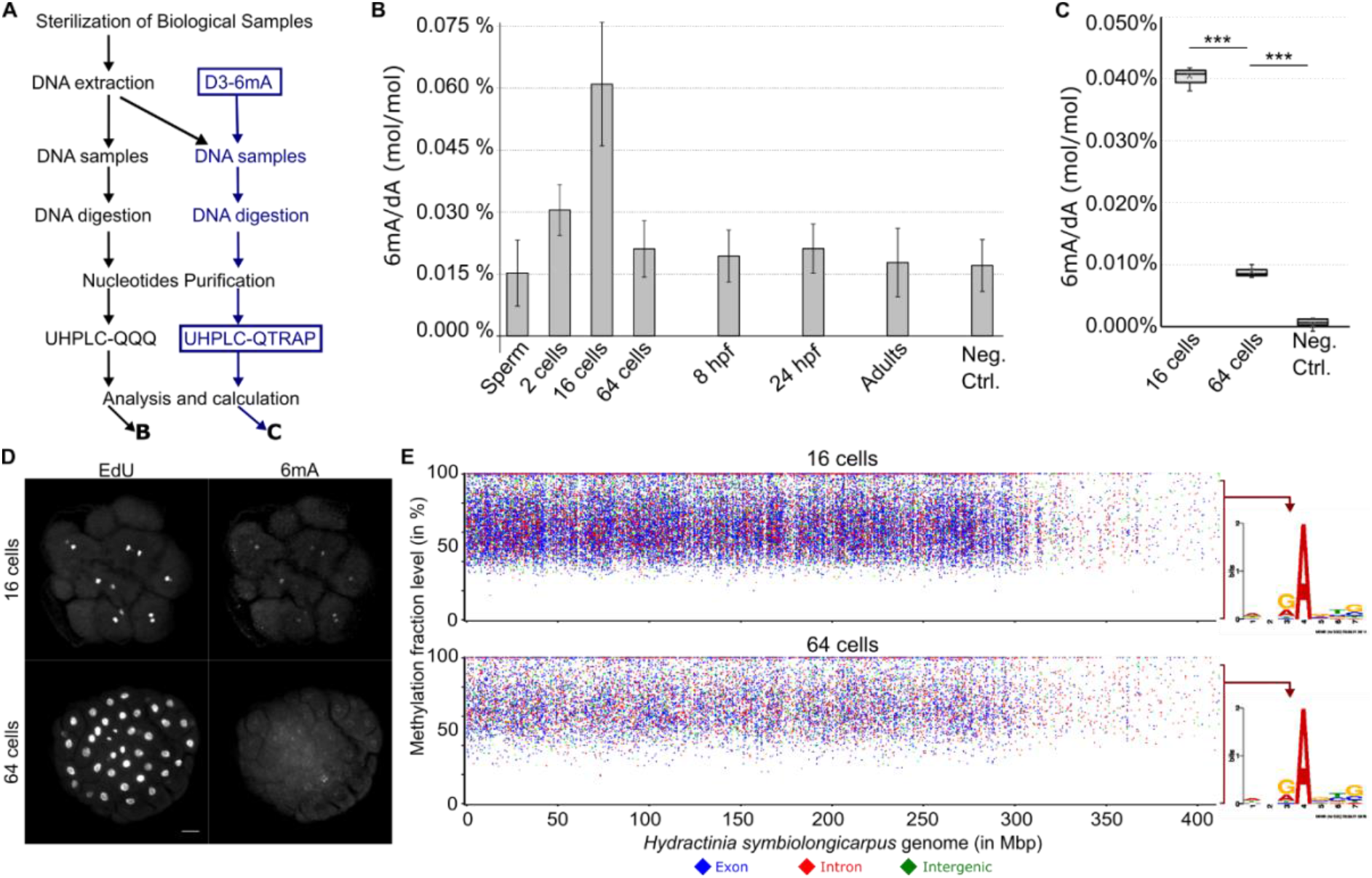
The dynamics and distribution of 6mA during *Hydractinia* early embryogenesis. **A.** Schematic of two independent methods to measure 6mA/dA levels: UHPLC-QQQ and UHPLC-QTRAP with D3-6mA internal standard (in blue). **B**. Levels of 6mA/dA (mol/mol) from seven stages of *Hydractinia* development measured by UHPLC-QQQ, calculated using external standard curve. **C**. UHPLC-QTRAP detection of 6mA/dA levels of *Hydractinia* genome from 16- and 64-cells embryos. **D**. Whole-mount immunofluorescence of 6mA from 16- and 64-cell stages of *Hydractinia*. **E**. Distribution of A sites that were detected to be methylated in the genomes of 16- and 64-cell stage, plotted against the percentage of SMRT-seq reads that showed methylation at each site. Consensus sequences of 6mA sites where the methylation level is between 0-95% are shown right to the graph, indicating that no motif can be deduced.

To rule out the possibility of bacterial contamination with high amounts of 6mA, we used an anti-6mA antibody for immunofluorescence (IF) in fixed embryos. The 6mA signal was visible in nuclei of *Hydractinia* cells (Figure 1D & S1B) and could be abolished by DNase treatment, but not by RNase treatment (Figure S1C); this observation is consistent with methylation of the animal’s nuclear DNA.

Next, we performed single-molecule real-time sequencing (SMRT-Seq) to investigate the distribution of 6mA in the genome of 16- and 64-cell stage embryos and adults. The data of methylated A sites was filtered by a combination of interpulse duration (IPD) ratio >3.0, read count >10, and p-value <0.05 following a recently published guideline (Zhu et al., 2018). Overall, the numbers of methylated A-loci were consistent with the dynamics of the 6mA/A detected by UHPLC-QTRAP, being high at the 16-cell stage and low at the 64-cell stage (Figure S1D-E). However, over 90% of A-loci were found to be inconsistently methylated across SMRT-seq reads from any given developmental stage (16- and 64-cell embryos, and adults; Figure 1E, S1D & F), indicating heterogeneity in methylated A-loci across cells that are expected to be uniform, particularly at the 16-cell stage (Kraus et al., 2014). Only about 7% of the loci were methylated in 100% of the reads (Supplemental File 1), and only 532 of the loci that were methylated in over 95% were shared between the 16- and 64-cell stages (Figure S1G). Finally, no motif representing the sequence context of all 6mA loci could be generated (Figure 1E & S2). The motif generated from the 88 loci that were methylated in over 95% of the reads across all developmental stages examined was 5’-GACCG-3’ (Figure S1G). This motif does not include an ApT context, suggesting that 6mA is not heritable in *Hydractinia* (Figure S1G and S2). Based on the above data, we conclude that 6mA marks are randomly distributed in the embryonic genome.

### *Alkbh1* acts as a 6mA eraser in *Hydractinia* embryos

ALKBH1 has been reported to function as a 6mA demethylation initiator enzyme in animals (Tian et al., 2020; Wu *et al*., 2016). The *Hydractinia* genome encodes a single *Alkbh1* homolog (Figure S3) that we tested to deduce its potential role in 6mA clearance. For this, we designed a specific shRNA-targeting *Alkbh1* (*shAlkbh1*; Figure S4) and injected it into zygotes. Embryos injected with a shRNA-targeting *GFP* (*shGFP*) were used as a negative control (Figure 2A & S4). Confocal imaging of anti-6mA immunofluorescence in 64-cell embryos showed that, in *shAlkbh1* injected embryos, 6mA signals were higher when compared with those from *shGFP*-injected ones (Figure 2A). Co-injection of *shAlkbh1* and *Alkbh1* mRNA carrying four silent mutations (rendering it resistant to the *shAlkbh1*) partially rescued the 6mA signal (Figure 2A-B). To confirm these results, we electroporated *shAlkbh1* into zygotes, extracted genomic DNA at the 64-cell stage, and then analyzed the 6mA content by UHPLC-QTRAP mass spectrometry with [^3^D1]-6mA as internal standard. We found a significantly higher level of 6mA in *shAlkbh1* electroporated embryos as compared to *shGFP* electroporated ones at the 64-cell stage (Figure 2C), consistent with what was observed in the above-described IF studies. These results confirm that Alkbh1 acts in erasing 6mA from the genome of early *Hydractinia* embryos.

**Figure 2.**
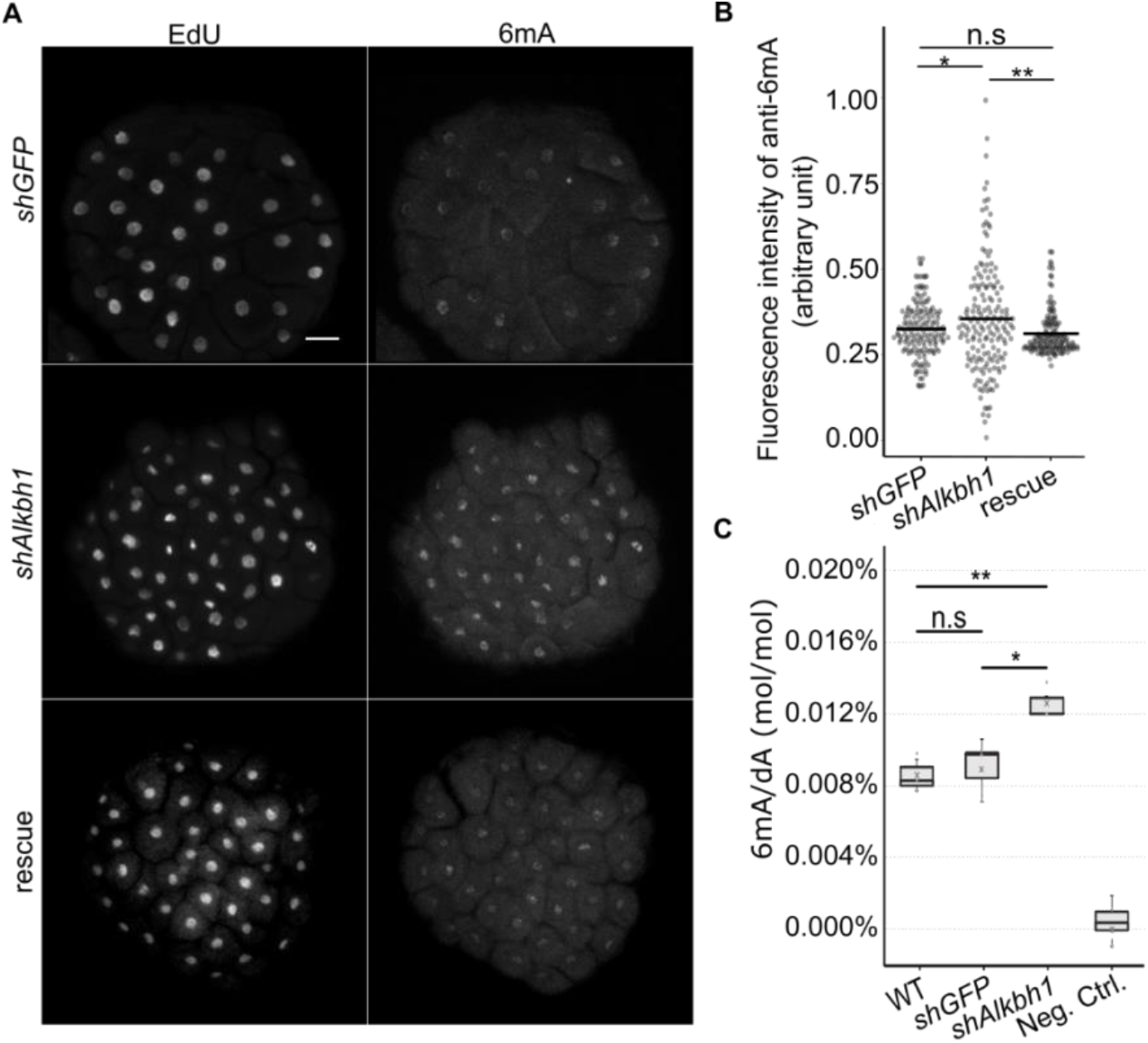
Alkbh1 removes genomic 6mA in *Hydractinia* embryos. **A**. Whole-mount immunofluorescence of anti-6mA in 64-128-cell embryos upon injection of *shGFP* (as control), *shAlkbh1*, and rescue (see text). **B**. Relative quantification of anti-6mA signals from immunofluorescence images (in triplicate). **C**. UHPLC-QTRAP quantification of *shAlkbh1*-electroporated embryos showing significantly higher level of 6mA/dA (P<0.05) compared to *shGFP* electroporated embryos and to wild type embryos at 64-128 cell stage. n.s: not significantly different (P=>0.05). * significantly different with P value < 0.05, ** significantly different with P value < 0.01.

### Zygotic genome activation follows 6mA clearance

In many animals, early embryos rely on maternal RNAs, activating their own genomes only at later developmental stages. Given the dynamic levels of 6mA in early embryos, we hypothesized that 6mA regulates the activation of the *Hydractinia* zygotic genome. To determine the stage at which zygotic transcription is activated, we used EU incorporation assays to visualize nascent RNA (Figure 3A) and established that a major transcriptional wave commences at the 64-cell stage, with little or no EU incorporation observed in earlier stages (Figure 3B-D). Therefore, it appears that a major wave of ZGA occurs immediately following the clearance of 6mA from the embryonic genome (Figure 1B-C & 3B).

**Figure 3.**
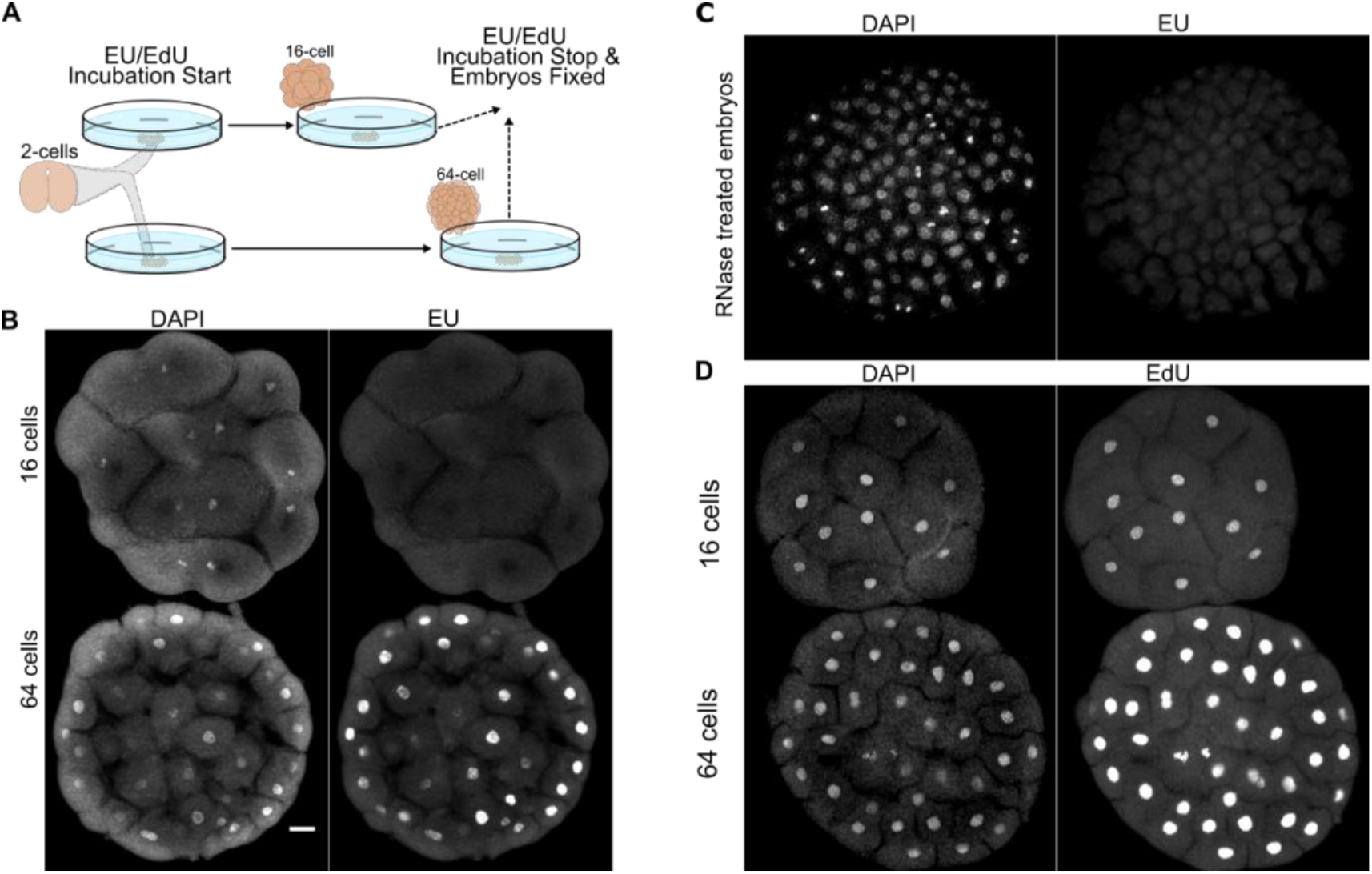
Zygotic Genome Activation at the 64-cell stage of *Hydractinia* embryos. **A**. EU/EdU incorporation experiment setup. **B**. High EU incorporation in 64-cell but undetectable in 16-cell embryos of *Hydractinia*. **C**. RNase treatment abolishes the EU signal. **D**. EdU is incorporated in 16- and 64-cell embryos. Scale bar: 20 μm.

### Alkbh1 knockdown delays zygotic genome activation

The occurrence of a major wave of ZGA immediately following 6mA clearance at the 64-cell stage prompted us to explore a possible functional link between these two phenomena. To examine this potential link, we injected *shAlkbh1* into zygotes to target *Alkbh1* mRNA and impede 6mA clearance. We then assessed zygotic transcription at the 64-cell stage by EU incorporation. We found that lowering Alkbh1 activity and the resulting elevated level of 6mA at the 64-cell stage (Figure 2) caused ZGA to be delayed by three cell cycles, commencing at the 512-cell stage instead of at the 64-cell stage as in untreated and *shGFP*-injected embryos (Figure 4 & S5). The late ZGA suggests that 6mA interferes with transcription, consistent with a previous study showing that genomic 6mA causes transcriptional pausing by stalling RNA polymerase II (Wang et al., 2017). The late recommencement of zygotic transcription in *Alkbh1-*knockdown embryos could have been enabled by 6mA dilution after DNA replication, assuming that 6mA incorporation was limited to occurring primarily in single-to 16-cell embryos. Delayed ZGA in *Alkbh1-*knockdown embryos caused no visible long-term defects; the embryos developed normally to planula larvae and successfully metamorphosed to primary polyps (Figure S5B).

**Figure 4.**
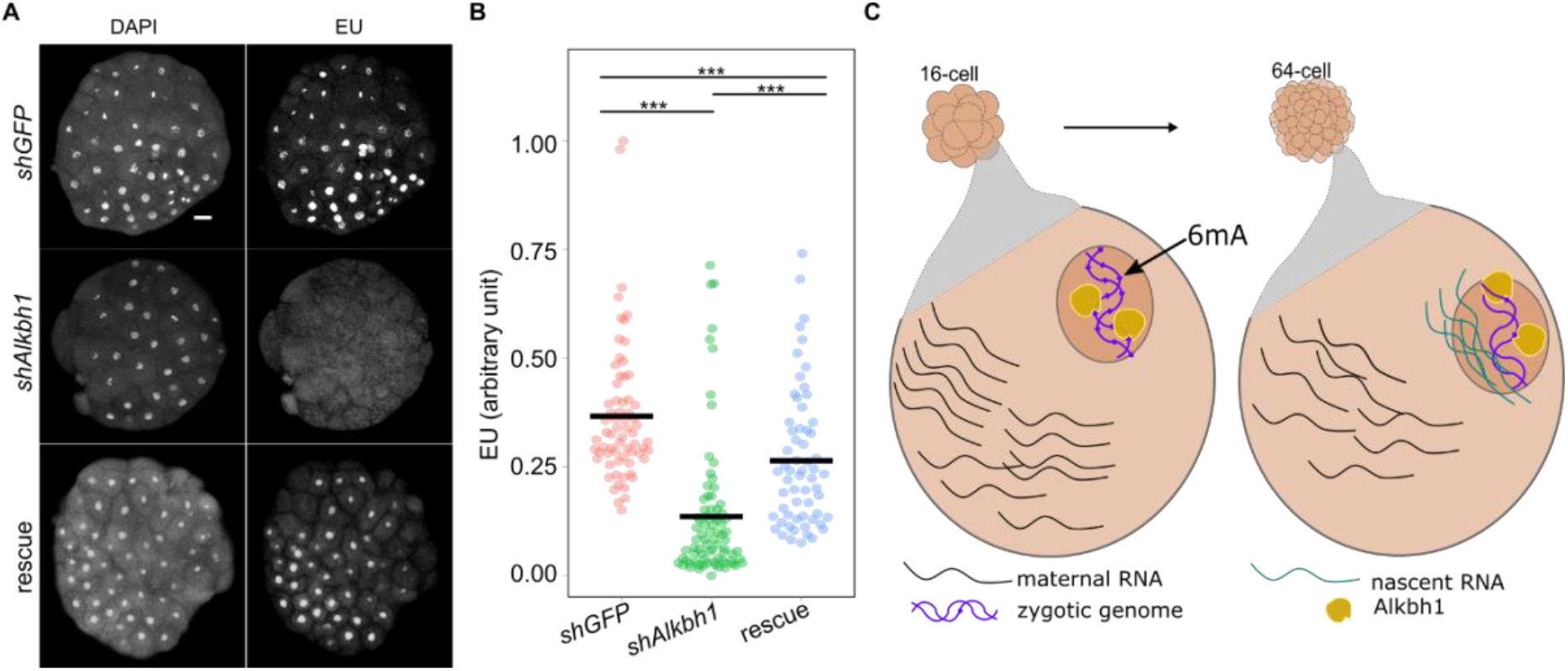
Knockdown of *Alkbh1* delays zygotic genome activation in *Hydractinia*. **A**. Whole-mount image of EU incorporation signals at 64 cells upon injection with *shGFP*, *shAlkbh1*, and rescue solution (see text). **B**. Relative quantification of EU signals (in triplicate). **C**. Model displaying the genomic 6mA removals by Alkbh1 prior to zygotic genome activation.

### The source of 6mA in the embryonic genome

To address how 6mA is incorporated into the *Hydractinia* genome between the 2- and 16-cell stages, we initially focused on *Mettl4 and N6amt1*, homologs of both of which have been proposed to function as 6mA methyltransferases in other animals (Greer *et al*., 2015; Xiao *et al*., 2018). The *Hydractinia* genome encodes one copy of each of the genes (Figure S6). Of note, *Hydractinia* and other animals’ N6AMT1 proteins contain no clear nuclear localization signal (Table S1). The likely inability of N6amt1 to act on nuclear DNA is inconsistent with a role as 6mA methyltransferases. If one of these genes (*N6amt1* or *Mettl4*) functioned as a 6mA methyltransferase, their downregulation would be expected to cause premature ZGA due to the absence of 6mA (Figure S7A). However, downregulation of both genes using shRNA did not result in premature ZGA (Figure S7B). Consistent with our results, recent reports show that Mettl4 and N6amt1 do not deposit 6mA in mammalian cells (Liu et al., 2020; Xie *et al*., 2018).

## DISCUSSION

A possible alternative source for methylated adenosine is m6A-marked RNA. In animals, maternal transcripts are degraded prior to ZGA (Chen et al., 2019; Varnum and Wormington, 1990), with m6A acting as a degradation mark (Ivanova et al., 2017; Zhao et al., 2017). We propose that methylated adenine from degraded maternal RNA is recycled through the salvage pathway and fuels methylated DNA synthesis during *Hydractinia* embryonic cleavage. Five observations are consistent with this hypothesis. First, we have performed HPLC-MS/MS experiments and find that m6A-marked RNAs are indeed degraded between the 2-cell and the 16-cell stages in *Hydractinia* embryos (Figure 5A), providing high amounts of methylated adenosine. Second, continuous RNR inhibition by hydroxyurea, starting with zygotes, stalled replication at the 8-cell stage (Figure 5B), indicating the depletion of maternally provided dNTPs and the requirement for NTP-dNTP conversion prior to this stage. Third, the random distribution of 6mA in the genome (Figure 1E) suggests a non-selective incorporation of 6mA into replicating DNA. Fourth, the delayed ZGA upon *Alkbh1* knockdown (Figure 3B and S6A) and the lack of premature ZGA following *N6amt1*/*Mettl4* knockdown (Figure S7B) indicate a lack of methyltransferase that maintains 6mA through embryogenesis. Finally, labeling gravid females with EU, followed by spawning and fertilization, resulted in embryos that had the signal in their nuclei (Figure S8). This is consistent with studies done in mammalian cells, showing that m6A ribonucleotides can be converted to 6mA deoxynucleotides and incorporated into the genome through a metabolic pathway that is conserved in animals (Musheev et al., 2020).

**Figure 5.**
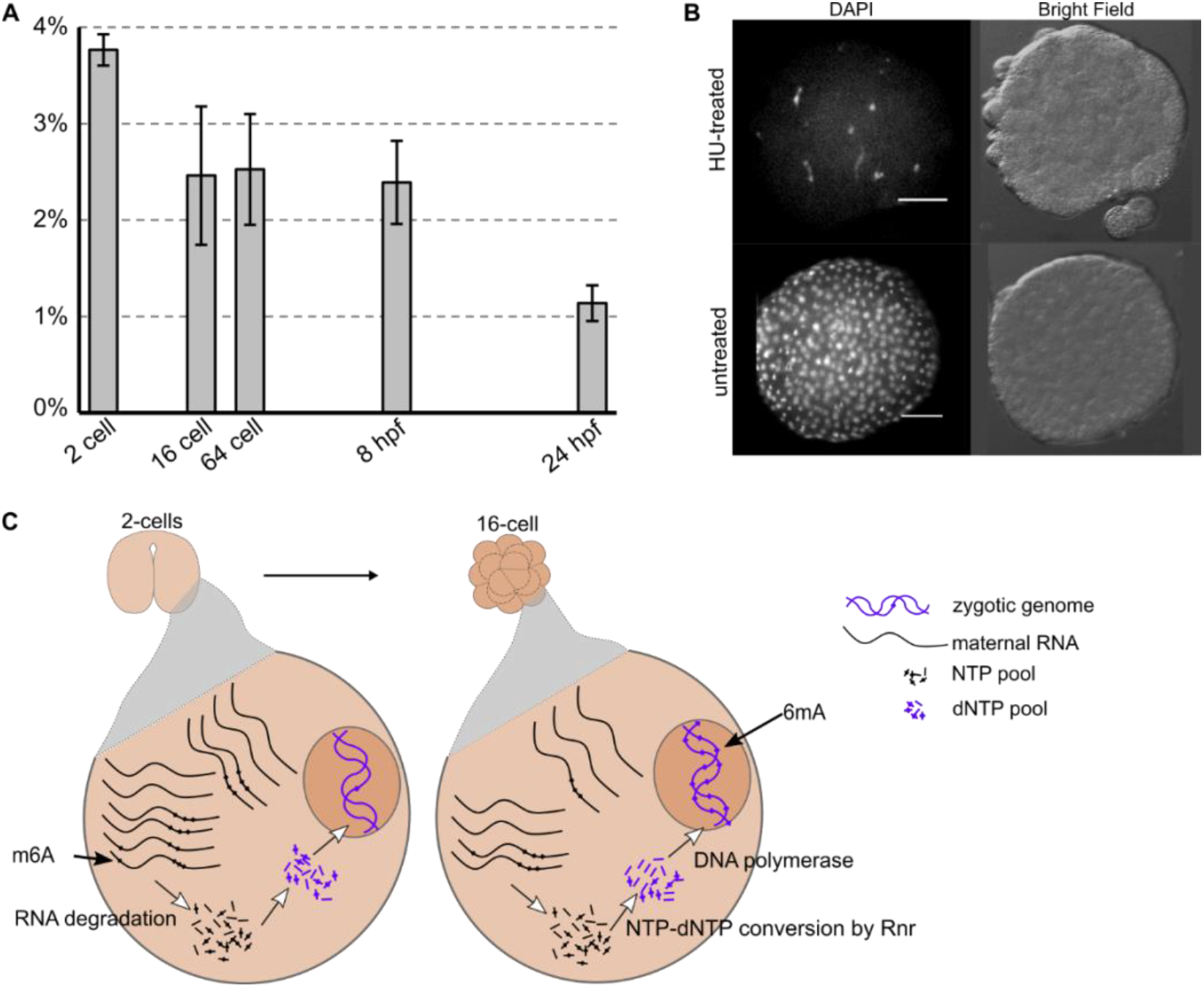
The maternal RNA recycling hypothesis and the evidence supporting it. **A**. Rapid decline of m6A-marked maternal RNA occurs between the 2- to 16-cell stages, analyzed by UHPLC-QQQ of m6A/A (mol/mol) from five *Hydractinia* developmental stages. **B**. Replication stall at 8-16 nuclei following hydroxyurea treatment. The control shows a normal number of nuclei. **C**. Model of stepwise m6A to 6mA conversion, followed by genomic incorporation of 6mA during early embryogenesis of *Hydractinia*.

An inverse correlation between zygotic transcription and 6mA during early embryogenesis can be inferred from studies using zebrafish and *Drosophila* (reviewed in ref (Bochtler and Fernandes, 2020)). Therefore, the model we propose for *Hydractinia* (Figure 5c) may be a general characteristic of all animals. Taken together, we conclude that 6mA is randomly and passively accumulated within the *Hydractinia* genome. This leads to the inhibition of transcription, particularly in early embryos, but is not epigenetic in nature. Alkbh1 is essentially a ‘cleaner’, keeping the genome 6mA-free and transcriptionally active.

## Acknowledgments

We thank our laboratory members for support and discussions, the NIH Intramural Sequencing Center (NISC) for generating the sequence data, and Jonathan J. Henry (University of Illinois at Urbana-Champaign) for advice on electroporation. Markus Müller (University of Munich) is kindly acknowledged for providing D3-6mA and for comments. Confocal images were taken at the Centre for Microscopy and Imaging Core Facility at NUI Galway. HPLC-MS/MS data were obtained at the Mass Spectrometry Facility at NUI Galway. We would like to thank Paul Gonzalez at the National Human Genome Research Institute (NHGRI) of the National Institutes of Health (NIH) for his thoughtful insights and advice regarding computational approaches for data analyses. We also thank Anh-Dao Nguyen at NHGRI/NIH for her efforts in making the data generated in the course of this study publicly available through the Hydractinia Genome Project Portal (https://research.nhgri.nih.gov/hydractinia).

## Author contributions

F, SGG, and UF initiated and conceptualized the project. F performed all laboratory experiments and analyzed data. MSS established electroporation. SNB, CES, SGG, ADB, and F performed the computational analyses. F and UF wrote the manuscript.

## Data availability

The data generated in the course of this study are publicly available through the *Hydractinia* Genome Project Portal (https://research.nhgri.nih.gov/hydractinia). Corresponding data is archived in the NCBI Sequence Read Archive (SRA) under BioProject PRJNA807936.

---

**Figure S1.**
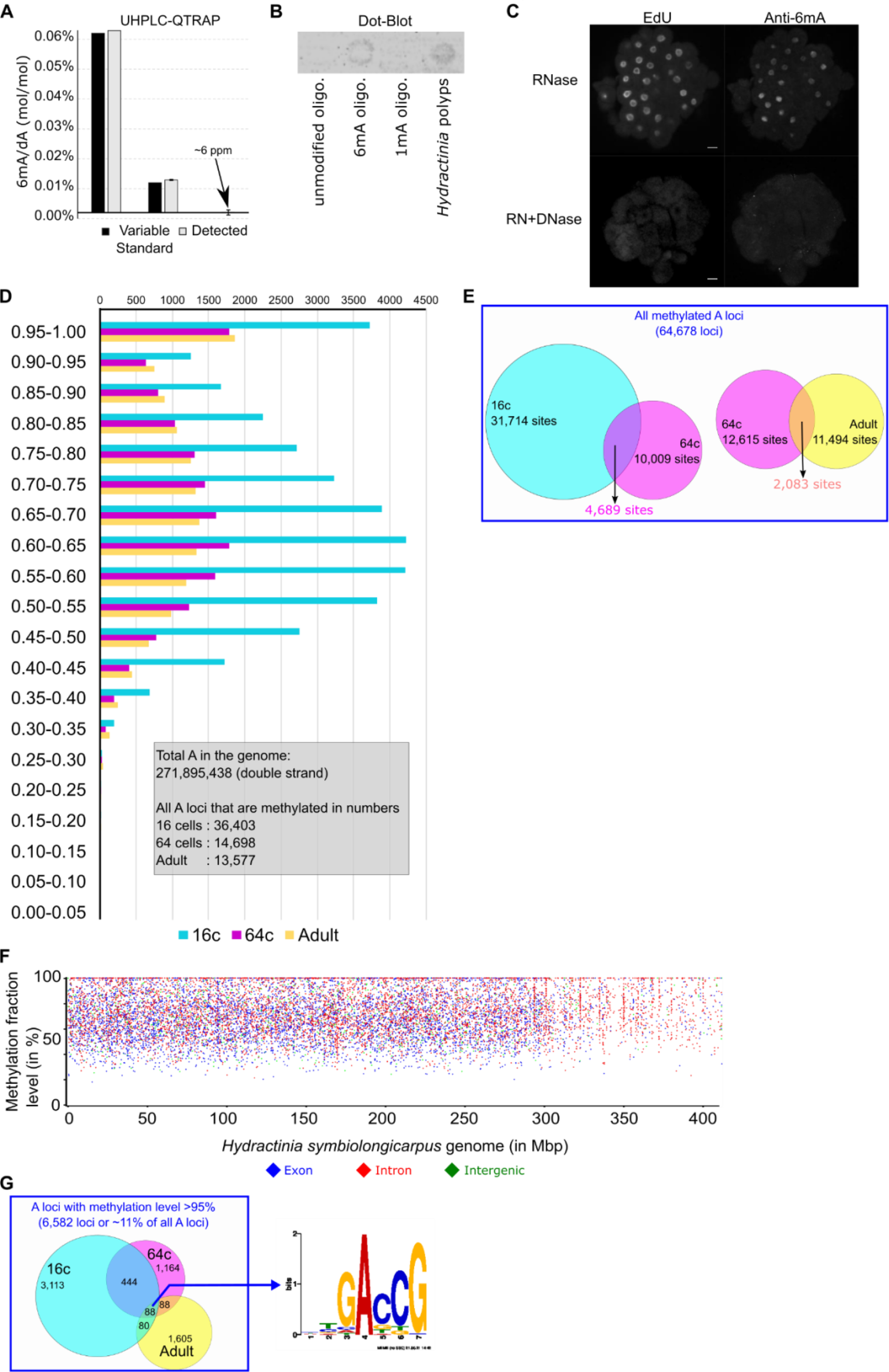
Detection and distribution of 6mA in the genome of *Hydractinia symbiolongicarpus*. **A**. Detection of 6mA in reference solutions (0%, 0.01%, 0.06%) by UHPLC-QTRAP. **B**. Anti-6mA specificity assay by dot-blot. Each spot contains 200 ng DNA. 6mA/1mA oligos were prepared at 0.1% of modified-A/dA. **C**. DNase but not RNase treatment can abolish the signal of anti-6mA immunofluorescence. Scale bars: 20 μm. **D**. Methylation level distribution on 16c, 64c, and adult genomes of *Hydractinia*. **E**. Venn diagram displaying the overlapping methylated A sites between 16c and 64c and between 64c and adult genomes. **F**. Distribution of A sites that were detected to be methylated in the genomes of adult specimens, plotted against the percentage of SMRT-seq reads that showed methylation at each site. **G**. Venn Diagram displaying the overlapping A sites between three genome that are always methylated and the consensus sequence generated by MEME-Chip of the 88 overlapping methylated A loci.

**Figure S2.**
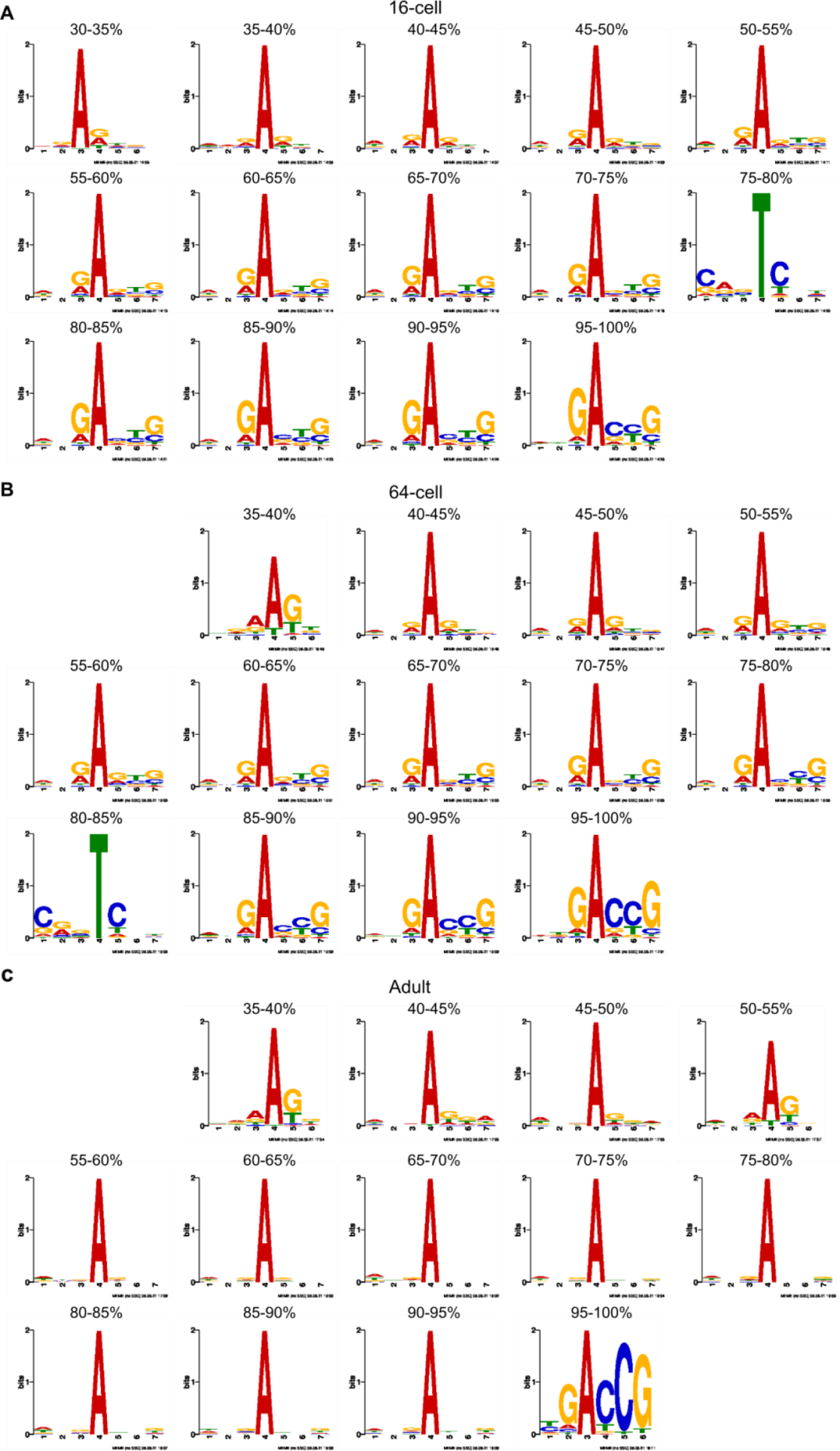
The consensus sequence generated by MEME-Chip of the methylated A loci in their respective methylation fraction. **A**. 16-cell. **B**. 64-cell. **C**. Adult.

**Figure S3.**
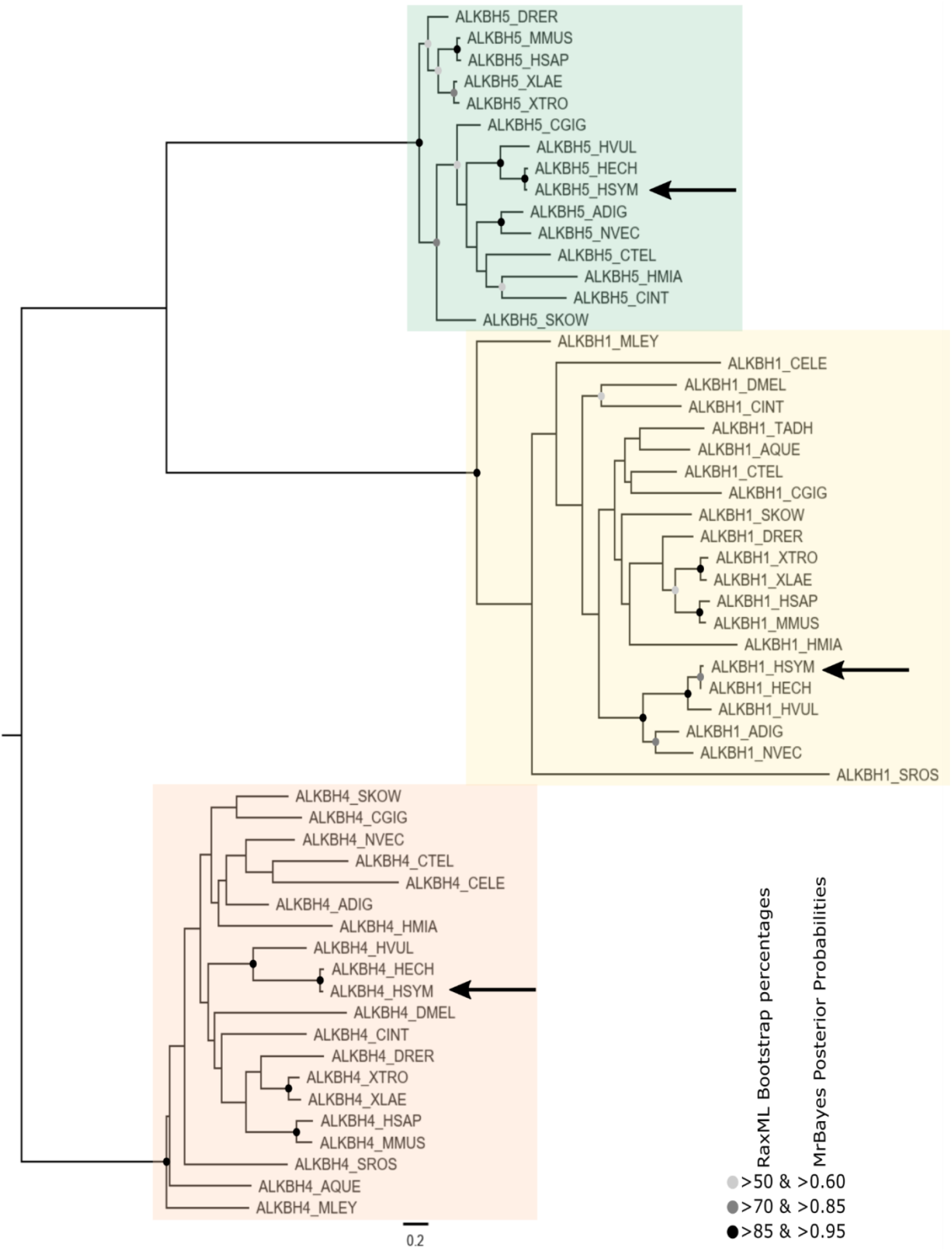
Phylogenetic analysis of Alkbh proteins. Maximum likelihood phylogenetic tree. Nodes supported by maximum likelihood bootstrap percentage and Bayesian inference posterior probability values are marked with greyscale circles as annotated. Alkbh homologs of *Hydractinia* are pointed by arrows. The abbreviation of the species are described in Table S2. The raw alignment data and fasta file of all the sequence used in this phylogeny are provided in Supplemental File 2.

**Figure S4.**
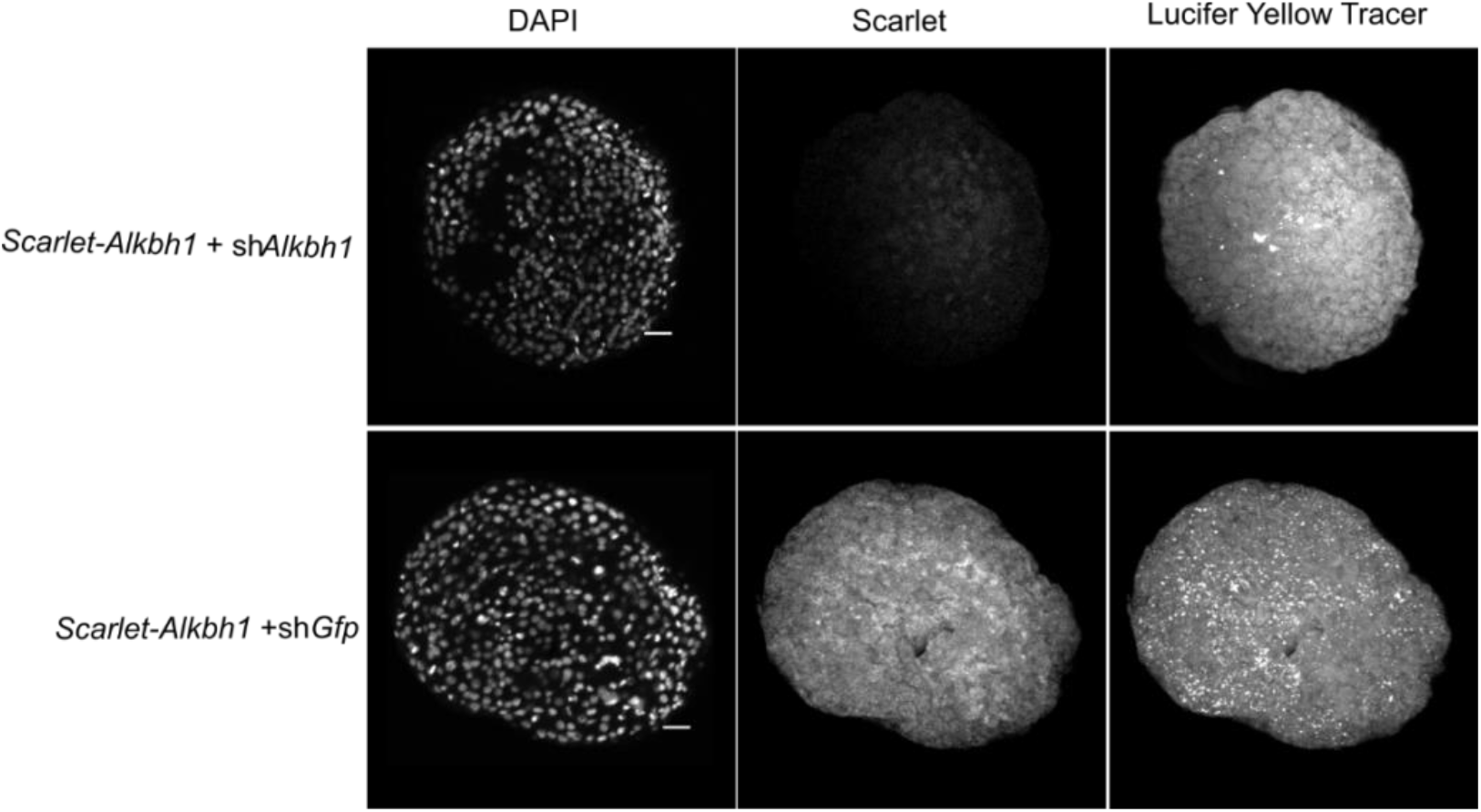
Synthetic mRNA encoding mScarlet fluorescence protein followed by the target sequence of *Hydractinia* sh*Alkbh1* were co-injected (1 μg/μl) with sh*Alkbh1* and *shGFP* (each 500 ng/μl). Strong signals of mScarlett in *shGFP* co-injection but not on *shAlkbh1* is indicative of successful knockdown effect by *shAlkbh1* at 15 hpf embryos. Scale bars: 20 μm.

**Figure S5.**
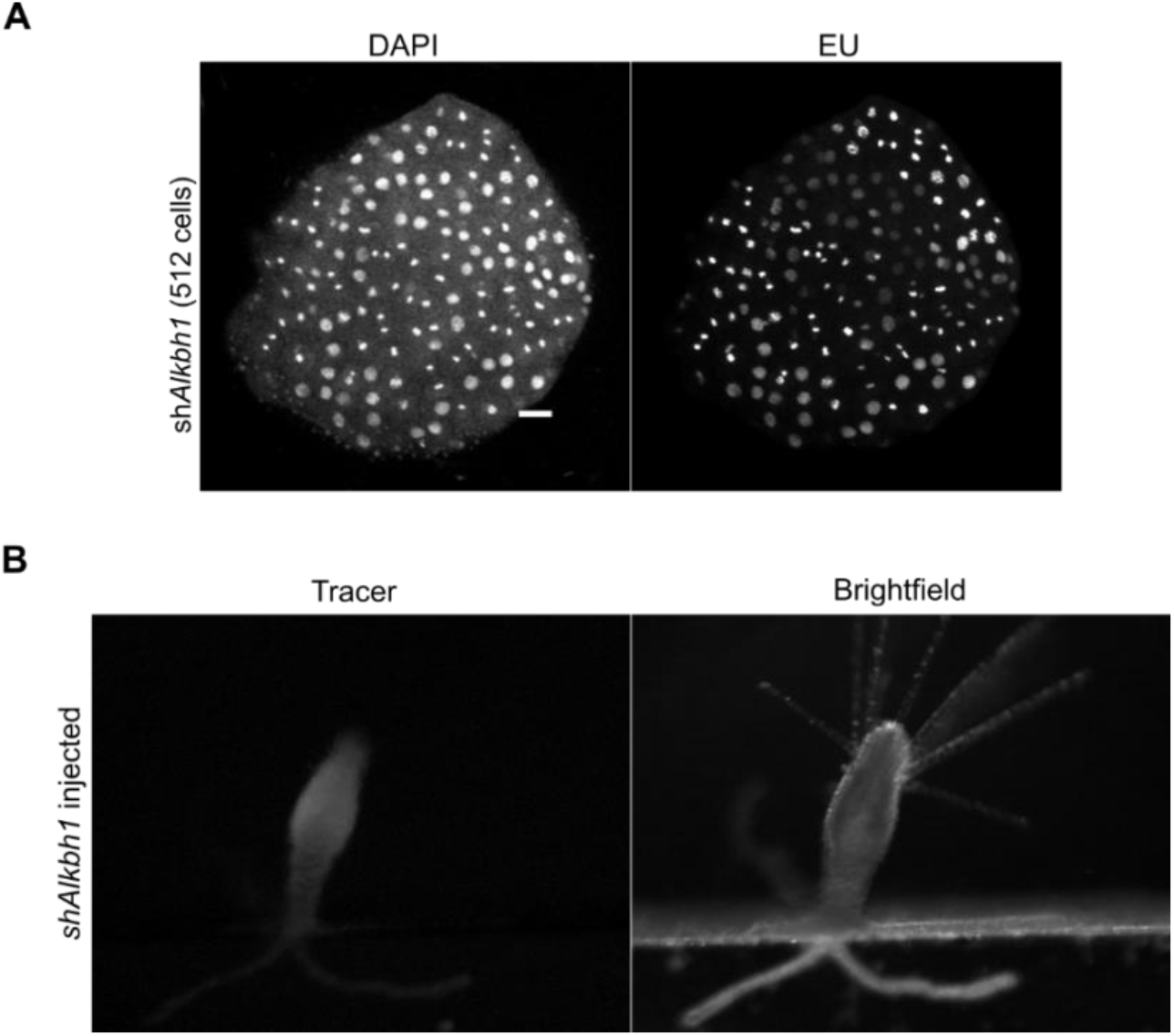
Knockdown of Alkbh1 delays ZGA but is not lethal. **A**. *Alkbh1* knockdown does not inhibit EU incorporation in 512-cell embryos. **B**. *shAlkbh1* injected embryo develops into a normal polyp. Scale bars: 20 μm.

**Figure S6.**
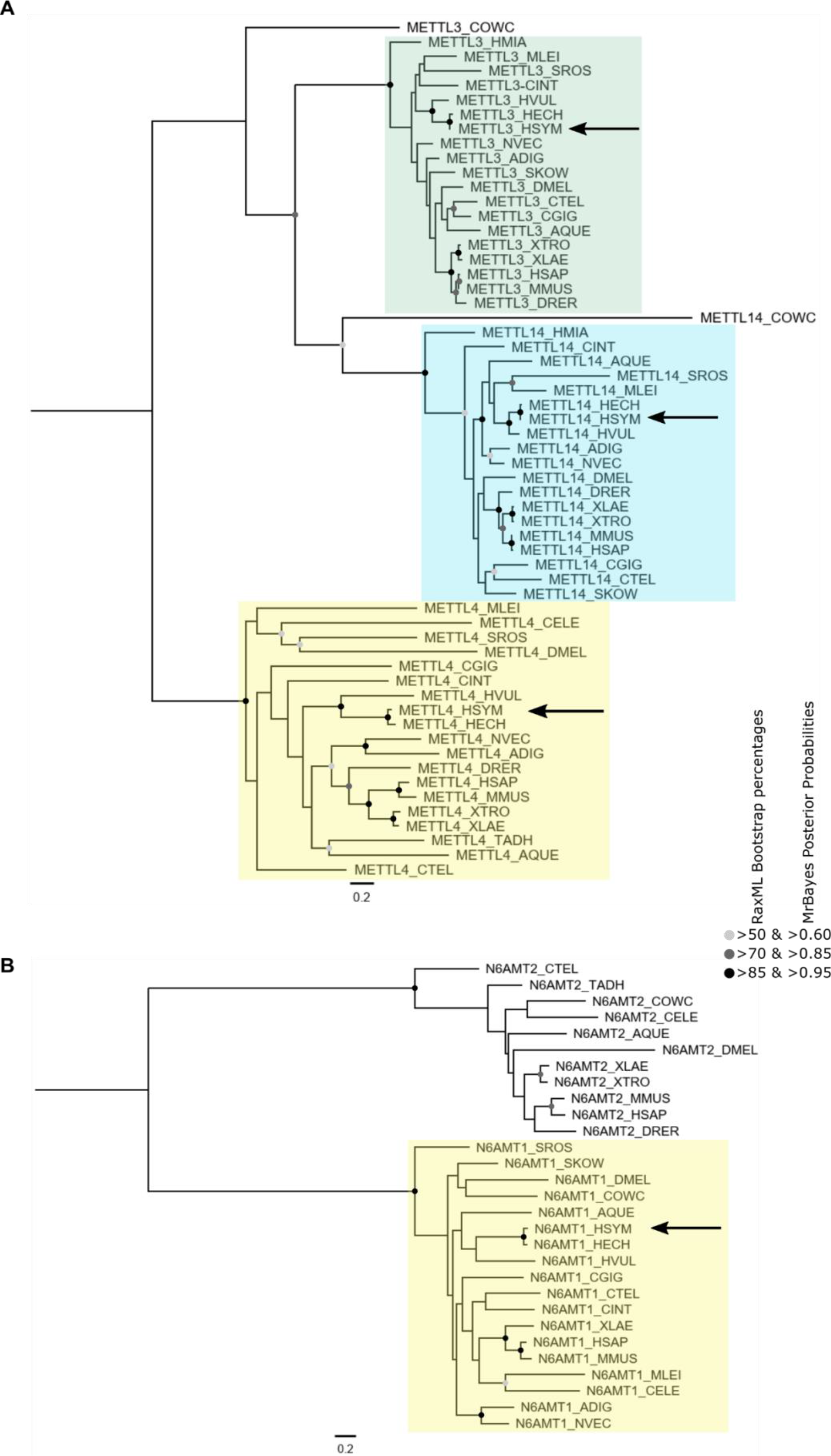
Phylogenetic analysis of Mettl and N6amt proteins. The trees represent a maximum likelihood phylogenety. The nodes with strong supports from maximum likelihood bootstrap percentages and Bayesian inference posterior probability are marked with a greyscale circle as annotated. Mettl4 and N6amt1 homologs of *Hydractinia* are pointed with arrows. The abbreviation of the species are described in Table S2. The raw alignment data and fasta file of all the sequence used in this phylogeny provided in Supplemental File 2.

**Figure S7.**
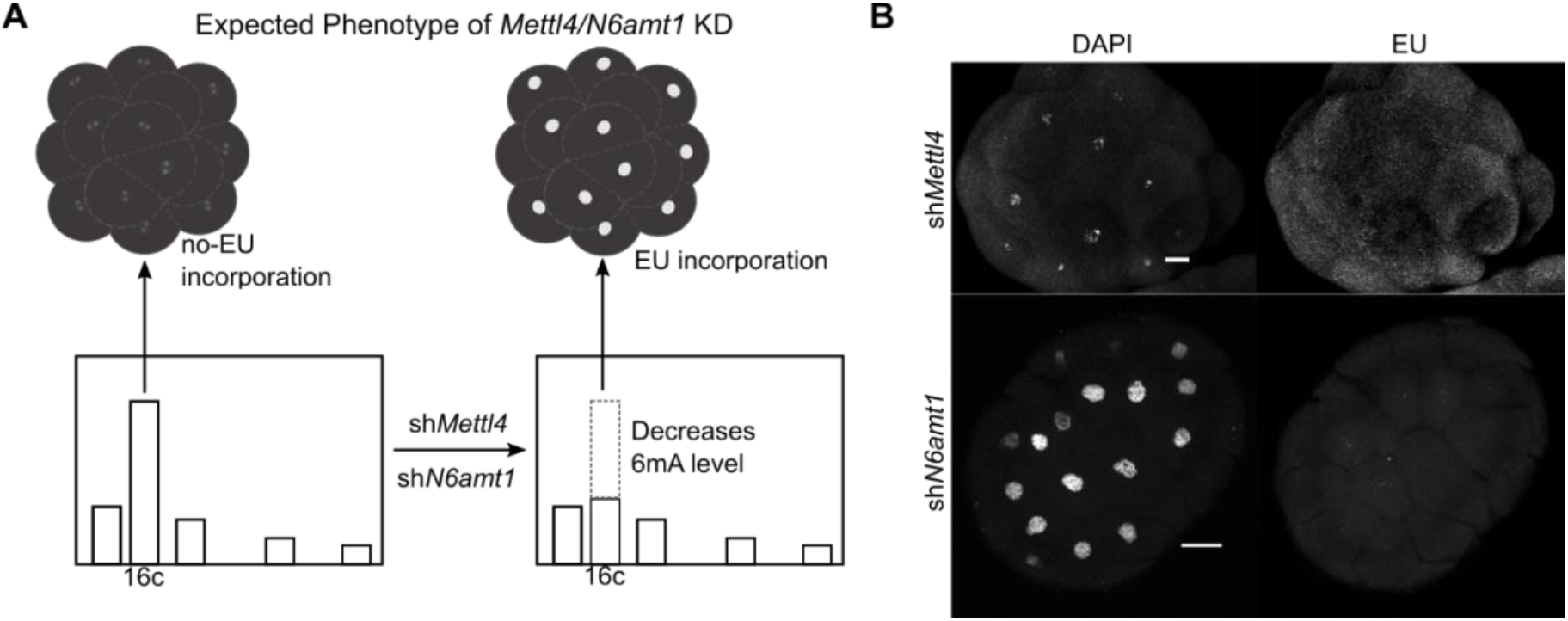
Knockdown of 6mA methyltransferase candidate do not display premature ZGA. **A**. Experiment setup. Knockdown of *Metll4*/*N6amt1* would be expected to result in premature ZGA if these enzymes were acting as 6mA methyltransferases. **B**. *Mettl4* and *N6amt1* knockdown does result in premature ZGA, suggesting that they do not act as 6mA methyltransferases. Scale bars: 20 μm.

**Figure S8.**
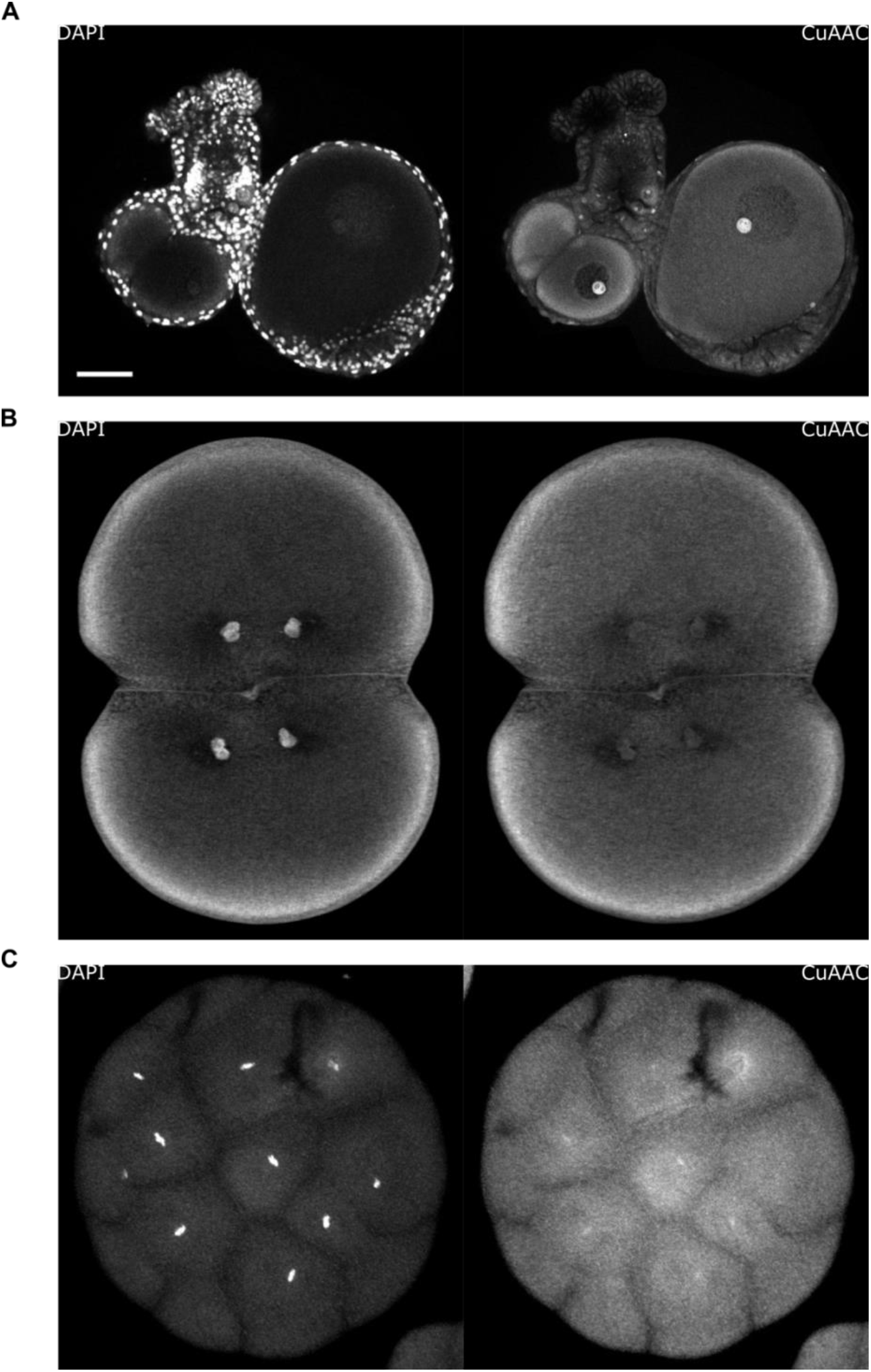
Transfer of nucleotides from maternal RNA to zygotic DNA. **A**. EU incorporation into nascent maternal RNA by a gravid female shown by CuAAC-Alexa 488 reaction in the cytosol and nucleolus of oocytes. **B**. Cytosolic maternal RNA at 2/4-cell stage embryo. **C**. CuAAC-Alexa 488 reaction stains the zygotic DNA in a 16-cell stage embryo. Scale bar in A: 50 μm.

**Table S1.** Nuclear localization signal prediction of methylation associated enzymes in *Hydractinia*

| Enzyme | cNLS Score | NLSdb ConfNuc | 1<sup>st</sup>/2<sup>nd</sup> PSORT | 1<sup>st</sup> PSORT Score | 2<sup>nd</sup> PSORT Score |
| --- | --- | --- | --- | --- | --- |
| *DNMT3A_HSAP* | 18.5 | 6 | Cytop/Nucl | 16 | 10 |
| *Dnmt3_HSYM* | 7.5 | 3 | Nucl/Cytop-Nuc | 16.5 | 15.5 |
| *METTL4_HSAP* | 10.5 | 9 | Nucl/Cytop-Nuc | 20.5 | 15.5 |
| *Mettl4_HSYM* | 7.3 | 14 | Nucl/Cytop-Nuc | 19 | 16.5 |
| *Mettl4-CELE* | 10 | 0 | Nucl/Cytop-Nuc | 18 | 14 |
| *N6AMT1_HSAP* | 0 | 0 | Cytop/Cytop-Nuc | 17 | 15.5 |
| *N6amt1_HSYM* | 0 | 0 | Cytosk/Cytop | 15 | 7 |
| *ALKBH1_HSAP* | 6 | 0 | Cytop-Nuc/Cytop | 11.5 | 10.5 |
| *Alkbh1_HSYM* | 8.5 | 0 | Cytop/Cytop-Nuc | 23 | 17 |

The cNLS mapper score indicates the probability that a given protein sequence contains a NLS. 0 indicates no NLS detected. NLSdb ConfNuc indicates the number of protein sequence in the NLS database that match the enquiry. PSORT is the localization prediction made by Wolf-PSORT with 1<sup>st</sup>/2<sup>nd</sup> PSORT indicating the first and the second-best localization prediction, respectively. The 1<sup>st</sup> PSORT score indicates the number of proteins (with known localization) considered similar to the enquired protein within the 1<sup>st</sup> prediction. Cytop: Cytoplasmic, Nucl: Nuclear, Cytop-Nuc: Cytoplama-nuclear, Cytosk: cytoskeleton.

**Table S2.** List of species and their abbreviation used in phylogenetic trees

| **Represented Phyla** | **Species** | **Abbrev.** |
| --- | --- | --- |
| Choanoflagellata | *Salpingoeca rosetta* | SROS |
| Choanoflagellata | *Capsaspora owczarzaki* | COWC |
| Placozoa | *Trichoplax adhaerens* | TADH |
| Porifera | *Amphimedon queenslandica* | AQUE |
| Cnidaria: Anthozoa | *Nematostella vectensis* | NVEC |
| Cnidaria: Anthozoa | *Acropora digitifera* | ADIG |
| Cnidaria: Hydrozoa | *Hydra vulgaris* | HVUL |
| Cnidaria: Hydrozoa | *Hydractinia echinata* | HECH |
| Cnidaria: Hydrozoa | *Hydractinia symbiolongicarpus* | HSYM |
| Ctenophora | *Mnemiopsis leidyi* | MLEI |
| Xenacoelomorpha | *Hofstenia miamia* | HMIA |
| Ecdysozoa: Arthropoda | *Drosophila melanogaster* | DMEL |
| Ecdysozoa: Nematoda | *Caenorhabditis elegans* | CELE |
| Lophotrochozoa | *Capitella teleta* | CTEL |
| Lophotrochozoa | *Crassostrea gigas* | CGIG |
| Hemichordata | *Saccoglossus kowalevskii* | SKOW |
| Chordata: Tunicata | *Ciona intestinalis* | CINT |
| Chordata: Teleostei | *Danio rerio* | DRER |
| Chordata: Amphibia | *Xenopus laevis* | XLAE |
| Chordata: Amphibia | *Xenopus tropicalis* | XTRO |
| Chordata: Mammalia | *Mus musculus* | MMUS |
| Chordata: Mammalia | *Homo sapiens* | HSAP |

**Supplemental File 1. The methylation fractionation data of 6mA from IPD-analysis of PacBio reads of 16-cell, 64-cell and adult genome of *Hydractinia symbiolongicarpus* 291-10.**

**Supplemental File 2. The raw alignment data and fasta file of all the sequences used for molecular phylogeny of Alkbh, Mettl and N6amt.**

## MATERIALS AND METHOD

### Animal Husbandry and Embryos Collection

Clones of *Hydractinia*, male (291-10) and females (295-8, 295-6) strains, were grown as previously described (Frank et al., 2020). Zygotes were collected and immediately cleaned with sterile-filtered sea water. For manipulation and injection purposes, the zygotes were incubated in ice cold condition to delay cleavages.

### DNA Extraction

DNA was extracted from *Hydractinia* embryos and adult specimens using Phenol-Chloroform and glycogen precipitation protocols. Following RNaseA (ThermoScientific #EN0531) and RNaseT1 (ThermoScientific #EN0541) treatment, the DNA was further purified using a standard column-based purification protocol. The purified DNA was then assessed by UV-Vis spectrophotometer, Qubit dsDNA-BR (ThermoScientific # Q32850) and Qubit RNA-HS assay (ThermoScientific # Q32852). Only DNA solutions with undetected level of RNA by Qubit RNA-HS assay were used.

### UHPLC-QQQ and -QTRAP for Determination of 6mA Levels

A total of 2 μg of DNA was prepared for digestion. For UHPLC-QTRAP, one picomole of ^3^D1-6mA was added to the solutions as internal standard. External standards were prepared from serial dilution of modified oligonucleotide (5’-^**6m**^**A**TCGATCG-‘3) solutions; variable standard solutions were prepared from the calculated combination of the above modified oligonucleotide and an unmodified oligonucleotide (5’-GGGCAGTACACAGACTATGTTG-‘3) solutions. DNA solutions were then denatured at 100°C for 5 minutes, chilled in ice for 2 minutes and digested following a protocol described before (Greer et al., 2015). After centrifugal ultra-filtration (MW cut-off 3 KDa, Amicon, Millipore #UFC500396), the nucleotide solutions were assessed by Nanodrop and Qubit dsDNA-HS assay. The total amount of DNA is expected to be equal by Nanodrop measurement before and after digestion. QUBIT dsDNA-HS was used to confirm zero dsDNA in the solutions. The digested DNA solutions (samples and standards) were then injected in 2 μl of volume into an Agilent 1100 HPLC system coupled to a triple quadrupole (QQQ) 6460 mass spectrometer (Agilent Technologies Ltd, Cork, Ireland), or injected in 6 μl volume into and an Agilent 1260 HPLC system coupled to an SciEx 4500 QTrap. Analytes separation by liquid chromatography were carried out using reverse-phase Zorbax SB-C18 column (2.1 mm width x 50 mm length; 1.8 μm particles), flow rate 250 μl/min using mobile phase A (0.1% formic acid solutions in water) and mobile phase B (0.1% formic acid in acetonitrile). To detect the analytes, the QQQ and the QTRAP modes were set to positive electrospray ionization and selective multiple reaction monitoring (MRM). Nucleosides were identified using the nucleoside precursor (parent) ion to product (daughter) ion mass transitions; dC (228.1/112.1), dA (252.1/136.1), 6mA (266.1/150.1) and ^3^D1-6mA (269.1/153.1). Mol of dA and 6mA from the QQQ were interpolated from standard curve rendered from serial dilution of digested external standards. The mol 6mA from QTRAP were calculated following the previously reported guideline using the direct comparison to the ^3^D1-6mA internal standards (Traube et al., 2019). The 6mA/dA ratio was calculated as the mol of 6mA per total mol of deoxyadenosine (dA + 6mA).

### Dot-Blot

Dot-blotting was performed on 200 ng of RNA-free dsDNA solutions and standard solution from unmodified and modified oligonucleotides (0% and 0.1% 6mA/dA) as described (Greer *et al*., 2015) on Amersham Hybond-N+ membrane (GE #RPN119B) using anti-6mA antibody (Synaptic System #202003).

### EU Incorporation and CuAAC Reaction

Cleaned embryos were incubated in 1 mM EU (Jena Bioscience #CLK-N002) for 45 minutes before being fixed in PFA+Ac solution (paraformaldehyde 4% and 0.5% freshly added glacial acetic acid (Fernández and Fuentes, 2013)) on a rocker at room temperature for 1 hour. The embryos were then rinsed in 200 mM glycine for 15 minutes, then permeabilized by PTx (3×15 minutes). The embryos were then rinsed in 1 ml of 2 M HCl for 45 minutes to denature the DNA as antigen retrieval step. The HCl was washed and embryos were neutralized with 1 ml 100 mM Tris-HCl pH 8.0 for 2 x 15 minutes. The embryos were then rinsed in 1 ml block-i1 solution (3% BSA (MP Biomedicals #11444296) and 0.25% Triton-X (MP-Biomedicals #11471632) in 1x PBS) overnight at 4°C on a rocker, followed by CuAAC reaction.

#### CuAAC Reaction

Ethynyl groups in EU/EdU act as the alkyne, which can react with fluorophore tagged azide through The Cu(I)-catalyzed alkyne-azide chemistry (CuAAC) reaction (Presolski et al., 2011). The CuAAC solutions (Jena Bioscience #CLK-074) were prepared freshly (Alexafluor488-picolylazides 2 μM, CuSO4 1 mM, THPTA 5 mM, and Na-Ascorbate 100 mM, in Sodium Phosphate buffer).

Next, embryos in the block-i1 solution brought back to room temperature. The block-i1 solution was then replaced with 500 μl CuAAC solutions and incubated on the rocker for at least 45 minutes in the dark at room temperature followed by two PTx washes. The DNA was then stained with DAPI and the embryos mounted for imaging.

### Wholemount Immunofluorescence

Embryos were incubated in 10 μM EdU (Jena Bioscience # CLK-N001) ~45 minutes before fixed by incubation in PAGA-T (20% PEG 6000 (Sigma #81260), 4% Glycerol (Sigma #G5516), 2.5% Acetic Acid, 56% Ethanol in 100 mM Tris-HCl pH 6.0 (Invitrogen # 15568025) (Zanini et al., 2012)) for 1 hour at 4°C. The fixed embryos were then washed with 1:3 mixture of PAGA-T and PBS-Triton (PTx, 0.5% Triton-X in 1x PBS). Permeabilization was done by further washes the fixed embryos with PTx for 15 minutes on a rocker at room temperature for three times.

Samples were then treated with 1:50 RNase solution (Mixture of RNaseA, T1 and H. (20 mg/ml, 1000 U/μl, and 10 U/μl, respectively)) and/or DNase (2 U/μl, NEB #M0303) at 37°C overnight. After one PBS wash, the embryos were rinsed in 1 ml of HCl 2 M for 45 minutes to denature the DNA as antigen retrieval step. The HCl was washed and embryos were neutralized with 1 ml 100 mM Tris-HCl pH 8.0 for 2 x 15 minutes. The embryos were then rinsed in 1 ml block-i1 solution (3% BSA and 0.25% Triton-X in PBS) for 1.5 hours at room temperature on a rocker.

Next, the block-i1 solution was replaced with 500 μl CuAAC solutions (described above) then incubated on the rocker for at least 45 minutes in the dark and room temperature followed by two PTx washes. The fixed embryos were rinsed in 1 ml block-i1 solution (3% BSA in PTx) overnight at 4°C before replaced with 200 μl of the Rabbit anti-6mA antibody solutions (diluted 1:8000 in block-i1, Synaptic Systems #202003) for one hour at room temperature. Then, the fixed embryos were washed in 1x PBS for 2×15 minutes then rinsed in 400 μl block-i2 solution (5% goat serum (ThermoFisher #16210064) and 3% BSA in PTx) for 2 hours at room temperature. Then, embryos were soaked in anti-rabbit Alexafluor 594 antibody (1:2000 in block-i2) for 1 hour at room temperature. Next, the embryos were washed three times with PBS and mounted for confocal microscope imaging.

### Image Preparation and Quantification

The mounted embryos were imaged by a confocal laser scanning microscope (Olympus FV1000). Known positive control samples were used to calibrate the confocal setup against the negative control ones (replacing primary antibody solution with blocking solutions or replacing EU/EdU soaking steps with seawater only). Once balance between the two controls was achieved at particular setup, this setup was used when images taken from samples slides on the same day of image acquisition.

Images were imported to ImageJ software (Schneider et al., 2012). Nuclei were the region of interest (ROI), thus we used the threshold approach to select nuclear regions from the DAPI channel as the ROI. These ROIs were then used to measure the mean fluorescence intensity (MFI) and corrected to the background ROI following the standard quantitation method (Shihan et al., 2021).

To compare the images, we normalized all MFI of the images to be compared by defining the highest MFI in the population as 1 and the lowest MFI value as 0, thus normalized MFI value were calculated using the following equation:

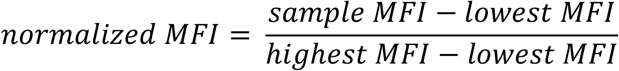

The normalized MFI was visualized using the online software at https://huygens.science.uva.nl/PlotsOfData/(Postma and Goedhart, 2019).

### SMRT-seq

Raw PacBio reads from adult polyps were provided by the NIH Intramural Sequencing Center (NISC) in fastq, bax.h5, and bash.h5 format. These files were converted to BAM format using bax2bam (SMRT Analysis; https://www.pacb.com/support/software-downloads/). Raw PacBio reads for 16-cell and 64-cell samples were provided in BAM format. BAM files for all three samples were aligned to the assembled genome with pbalign (https://github.com/PacificBiosciences/pbalign) in base modification identification mode, with the command-line version using default parameters and BAM formatted output). IpdSummary of SMRT Analysis (https://www.pacb.com/support/software-downloads/) was used to identify 6mA (using default options, with p-value 0.001, methyl fraction calculation, 6mA identification, and GFF output). The GFF output was then imported to Geneious for manual analysis. We achieved the recommended coverage (Zhu et al., 2018) in all datasets (16-cell, 64-cell, and adult polyps at 73x, 117x, and 120x, respectively).

Afterwards, 6mAs were filtered to remove those with IPD ratio below 3.0 (Zhu *et al*., 2018). Analysis of methylation motifs was performed with two different strategies. First, possible motifs were determined with MotifMaker using default options (SMRT Analysis; https://www.pacb.com/support/software-downloads/). To further confirm the lack of motif identification, all 6mA loci were separated into 20 groups based on their percent occurrence (in 5% intervals), and the regions 3 bp upstream and downstream of each 6mA were extracted. MEME-ChIP (Machanick and Bailey, 2011) was then used to identify consensus sequence in each group.

### RNA extraction and m6A Detection

Total RNAs was extracted from embryos of 2-4 cell, 16-32 cell, 64-128 cell stages, and 24 hours post-fertilization using TRIzol solution (ThermoScientific #15596026) followed by RNA binding onto columns (EpochLifeScience #1940) and on-column DNA digestion (Qiagen #79254). RNA was then eluted with nuclease free water, assessed with a Qubit RNA HS assay and electrophoresed along with RNA loading dyes (ThermoScientific #R0641) in denaturing formaldehyde agarose gel before visualization under UV illumination. High-quality RNA was then used detect 6mA using UHPLC-QQQ after RNase A/T1 overnight digestion and ultrafiltration with MRM of A (268.1/152.1) and m6A (282.1/166.1).

### Multiple Sequence Alignment (MSA) and Phylogenetic Tree Inferences

Sequences of Alkbh1 (Uniprot ID: P0CB42), N6AMT1 (Q9Y5N5), Alkbh4 (Q8MNT9), and Mettl4 (Q09956) were used as queries to retrieve orthologous sequences from a *Hydractinia symbiolongicarpus* transcriptome using tblastn. We retrieved the sequences of the respective homologs from each species from the uniprot database (www.uniprot.org) and Eensembl omics database (https://metazoa.ensembl.org/), which were imported into Geneious Prime 2019.0.4 software. We retrieved the homologous sequences of *Mnemiopsis leidyi* (NHGRI), *Hydra vulgaris* (NHGRI), *Hydractinia echinata* (NHGRI), *Saccoglossus kowalevskii* (OIST), and *Acropora digitifera* (OIST) from their specific respective database. Sequences were aligned in Geneious using MAFFT with the E-INS-i algorithm, a JTT PAM100 scoring matrix, and a gap penalty of 1.53 (Katoh and Standley, 2013).

The phylogenetic trees were built as a combination of three independent inferences from multiple sequence alignments. Firstly, a phylogenetic tree was built by RAxML 8.2.11 (Stamatakis, 2014) using the GAMMA LG protein model (default), rapid bootstrapping (10,000 replicates) and searching for best-scoring maximum likelihood tree algorithm. Secondly, a Bayesian phylogenetic tree was produced using MrBayes v.3.2.2 (Ronquist et al., 2012). The program was run using a fixed WAG substitution model (recommended by MrBayes trial with the respective MSA with 500 generations and sampled every 50^th^ generation) with gamma distributed rate variation across sites (“lset rates=gamma”) with four chains for 4 million generations. The run was sampled every 500^th^ generation and analysed with a 20% burn-in. These two methods of phylogenetic tree inference are available in Geneious. The consensus tree from maximum likelihood analysis was then exported and manually edited in InkScape to mark the nodes with support values as annotated from the two different methods of phylogenetic inference with greyscale dots.

### Localization Signal

Sequences from *Hydractinia symbiolongicarpus* and *Homo sapiens* homologous proteins were analysed for nuclear localisation signals by cNLS Mapper (Kosugi et al., 2009), by NLSdb (Bernhofer et al., 2018) and for protein sorting in general by Wolf Psort (Horton et al., 2007). The results retrieved and imported to Microsoft Excel for data visualization and presented as Table S2.

### *Alkbh1* knockdown and rescue experiment

Short-hairpin RNA were designed according to a previous report (DuBuc et al., 2020). T7 IVT kit was used to synthesize mRNA to confirm the efficacies of *shAlkbh1* by adding the endogenous target of *Alkbh1* sequences at the 3’ of *mScarlet* coding sequence. Rescue *Alkbh1* mRNA was designed by introducing four silent mutations, T861C, A864G, C865T, and A867G to render it unrecognizable by *shAlkbh1*.

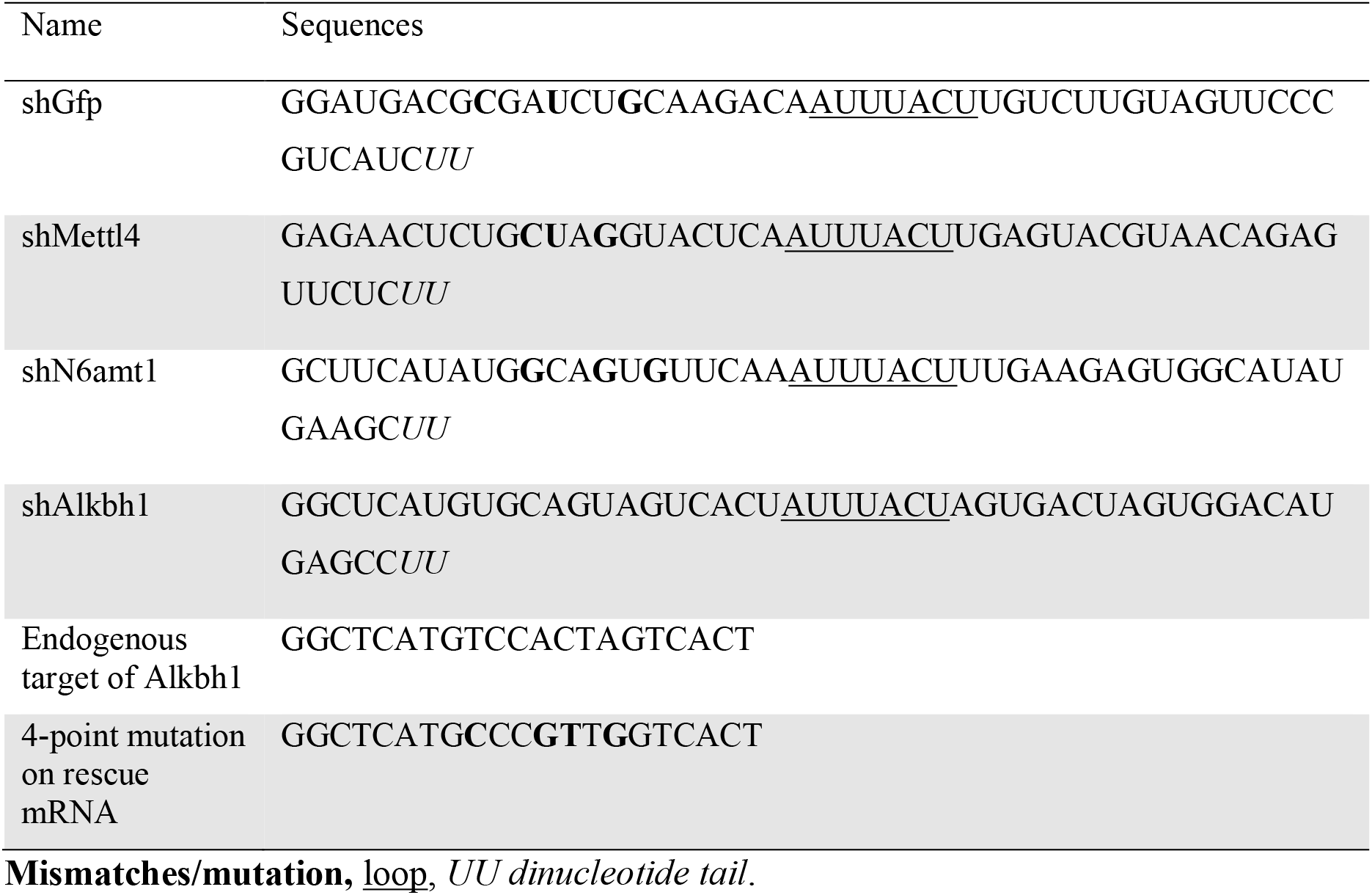
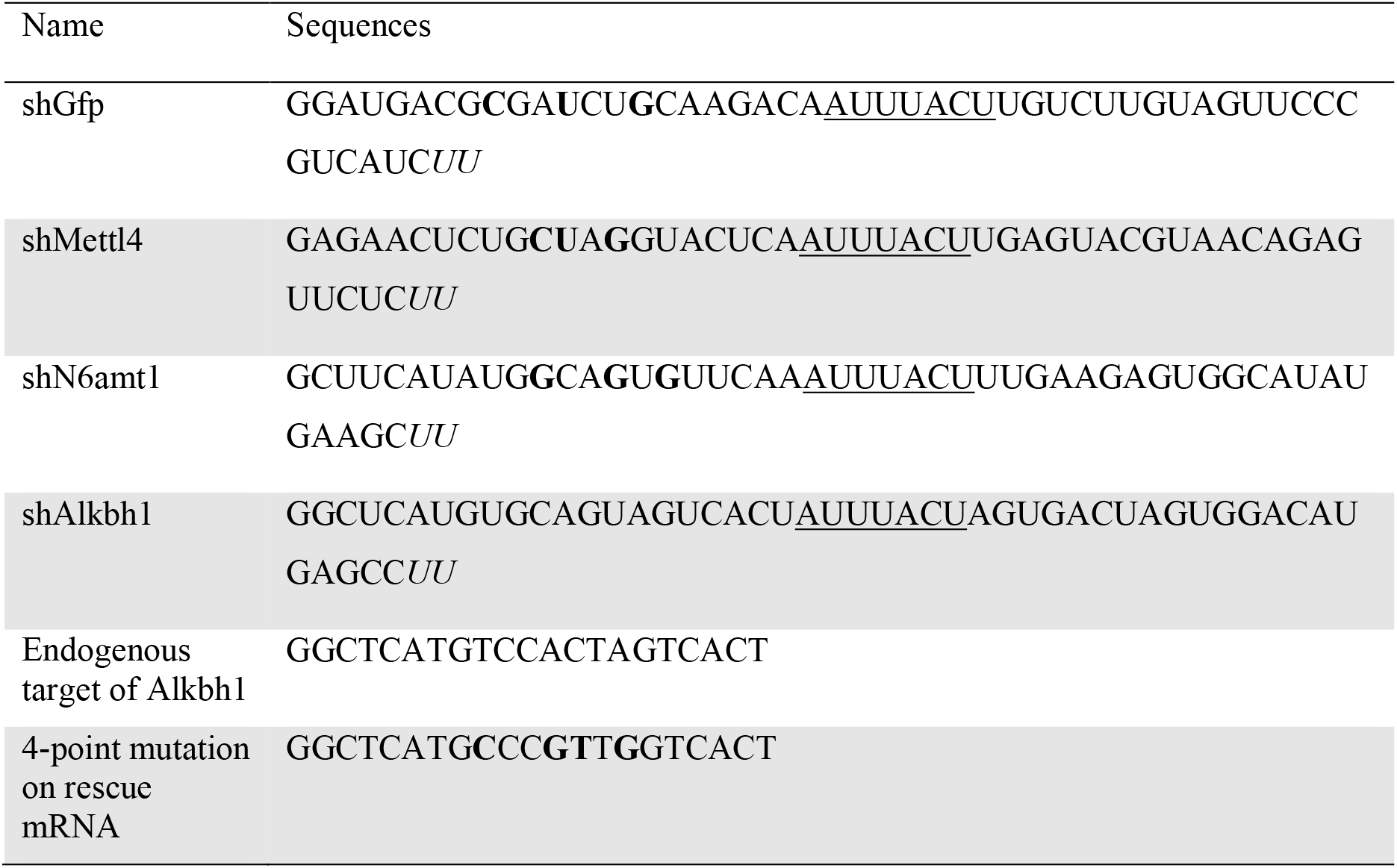

### Microinjection

Fertilized eggs were transferred to a Petri dish coated with 200-micron Nitex mesh screen. Zygotes are 180-200 microns and settled in the holes. Cells were injected, prior to first cleavage, using a Narishige IM 300 microinjection system. To delay cleavage, zygotes were stored on ice prior to injection.

### Electroporation

Zygotes were rigorously cleaned with filtered-sterile seawater then electroporated to insert *shAlkbh1* into the cell following the previously described protocol (Quiroga-Artigas et al., 2020) with Ficoll replaced by 1.54 M Mannitol. Next, zygotes were immediately transferred into a large volume of filtered-sterile seawater in glass Petri dish and left at room temperature for 1 hour before further cleaning and then used for DNA extraction, DNA digestion, and UHPLC-QTRAP protocols as described above.

### Hydroxyurea treatment

Cleaned 2-cell stage embryos were incubated in sea water with 10 mM Hydroxyurea (HU) and collected at the 256/512-cell stage, while the negative control embryos were incubated only in seawater. Both the HU-treated embryos and negative control were soaked in hoescht-33342 (diluted 1:2000 in seawater) for 15 minutes then mounted for image acquisition on an epifluorescence microscope.

